# Early acquisition of threat conditioning in a selectively-bred anxiety-like rat phenotype: regulation by maternal presence and FGF2

**DOI:** 10.1101/2021.04.30.442207

**Authors:** Amanda M. White, Da-Jeong Chang, Joanna H. Hider, Elaine K. Hebda-Bauer, Cortney A. Turner, Regina M. Sullivan, Huda Akil, Jacek Debiec

## Abstract

Temperament is an innate, stable predisposition towards particular emotional and behavioral responses. In humans, certain temperaments are associated with a heightened risk of developing anxiety later in life. Non-human animals, including rodents, also exhibit innate, stable dispositions; these are referred to as behavioral phenotypes. The interaction between behavioral phenotype and early life adverse events is critical for the development of maladaptive anxiety. Rodent studies of typically developing animals have identified a number of mechanisms that protect against aversive experiences in early life. One such mechanism is an early life quiescence of threat learning, which protects against the effects of stress and facilitates safety and attachment learning. However, little is known about the factors that alleviate the effects of early life aversive events on phenotypes vulnerable to pathological anxiety. Here, we examined threat learning and the stress response in selectively-bred infant rats that show an anxiety-like phenotype relative to typically developing animals. We investigated the potential roles of maternal presence and the anxiolytic neurotrophic factor fibroblast growth factor 2 (FGF2) in regulating threat learning and the stress response in infant anxiety-like phenotype animals. We observed that rats selectively-bred for anxiety-like behaviors could acquire conditioned freezing earlier in life than typically developing animals. FGF2 administration on postnatal day 1 (PND 1) and maternal presence during threat conditioning were both capable of suppressing this early emergence of conditioned freezing. However, neither FGF2 nor maternal presence during threat conditioning were associated with reduced corticosterone levels during threat conditioning. Our results suggest that although an anxiety-like phenotype may be associated with early threat learning, environmental factors (such as maternal presence) and pharmacological intervention (such as modulation of the FGF2 system) may be capable of counteracting that early aversive learning. Interventions in vulnerable infants may thus decrease the impact of aversive events.

## Introduction

Temperament is an innate, stable disposition that leads an individual to exhibit particular emotional and behavioral responses. Certain features of temperament are expressed very early in life and may predispose individuals for certain outcomes. For example, some children are comfortable in novel situations, while others display signs of discomfort and will cling to their parent or cease play in the presence of unfamiliar objects or individuals^1,2^. This inhibited response to novelty in early childhood appears to be stable and heritable and is referred to as behavioral inhibition^1–3^. Children who exhibit this behaviorally inhibited temperament are at a heightened risk for anxiety disorders^4^. Behavioral phenotypes characterized by inhibited behavior in novel environments can also be observed in other animal species, including non-human primates^5,6^ and rodents^7,8^.

After many generations of selectively breeding rats based on their response to a novel environment, the Akil lab has developed two lines of Sprague-Dawley (SD) rats that represent the extremes of this phenotype – those that show high activity in a novel environment and those that show low activity in a novel environment, which the authors term “Bred Low-Responders (bLR).” In addition to exhibiting differences in their exploratory behaviors, bLR rats tend to show increased anxiety-like behaviors as adults. Specifically, comparing to animals showing high activity in novel environments, bLR rats spend less time in the open arms of the elevated plus maze, spend less time in the light area of the light-dark box, and spend less time in the center of an open field^9^. Additionally, adult bLR rats show deficits in extinction of threat learning and extinction retention^10^.

Rodent studies using both genetically engineered and selectively bred models have identified a number of candidate neurobiological mechanisms underlying anxious-like phenotypes^11–14^. Several studies have pointed to the role of neurotrophic factors in the regulation of affective-related behaviors^15,16^. One of the neurotrophic factors involved in anxiety-like behavior in rats is fibroblast growth factor 2 (FGF2)^17^. In typically developing adult rats, lower endogenous levels of FGF2 have been associated with greater anxiety-like behavior in the elevated plus maze^18^ and higher cue-induced freezing^19^. Furthermore, knockdown or knock-out of FGF2 in typically developing rodents led to increased anxiety-like behaviors in the elevated plus maze^18,20^. Knock-out of FGF2 led to elevated baseline corticosterone levels, and enhanced corticosterone response to stress^20^. The FGF2 system also appears to play a critical role in the behavioral phenotype exhibited by bLR rats, as adult bLR rats show lower endogenous levels of FGF2 mRNA in the hippocampus^21^.

Early life intervention could modify some features of anxiety-like phenotypes, such as responses to stress and adversity. In typically developing animals, administration of FGF2 has been shown to influence threat learning, extinction, and anxiety-like behaviors^18,22–26^. In bLR rats, a single subcutaneous (s.c.) injection of FGF2 on the day after birth was associated with less anxiety-like behavior in adulthood than bLR rats given a vehicle injection^21^. Additionally, bLR rats that received an FGF2 injection showed improved extinction of conditioned threat responses and improved extinction retention in adulthood^10^.

The bLR model is particularly useful as it can shed light on the behavioral phenotypes that predict susceptibility to anxiety disorders and on the developmental time points at which preventive interventions may prove most effective. This is a critical question for translational applications, as anxiety disorders typically develop in childhood and early intervention may lead to improved outcomes later in life. However, not all children with behavioral inhibition will go on to develop anxiety disorders; the interaction between behavioral phenotype and the environment is critical for the development of maladaptive anxiety. One of the known experiential risk factors for anxiety disorders is an aversive experience. Most of what is known about the neurobiology of aversive experiences comes from human and animal studies of threat learning in phenotypically normal adults, and little is known about early life aversive experiences in the anxious-like phenotypes.

Studies using outbred rat pups show that until the animals start leaving the nest around postnatal day (PND) 10, the hypothalamic-pituitary-adrenal (HPA) axis^27,28^ and amygdala are generally quiescent, and Pavlovian threat learning does not typically occur^29,30^. It is hypothesized that the lower activity of the HPA axis prior to PND 10 is primarily responsible for the quiescence of threat learning at this age, as injection of corticosterone prior to threat conditioning in pups younger than PND 10 can drive synaptic plasticity in the amygdala and support the acquisition of threat learning until pups are PND 16 ^31–33^. Additionally, a dam that expresses defensive responses in the presence of a threat conditioned cue can acutely elevate corticosterone levels in pups and transfer that conditioned threat to her PND 6-7 pups^34^. Once rat pups reach ten days of age the HPA axis becomes more reactive to stressors^28^ and Pavlovian threat learning to olfactory cues emerges^29^. However, if the pup undergoes threat conditioning in the presence of a calm mother, threat learning and the accompanying corticosterone response is suppressed^32^. The low reactivity of the HPA axis in very young pups and the ability of maternal presence to extend the period of this low reactivity in pups from PND 10-15 are thought to be adaptive mechanisms by which the still-developing pup is protected from the deleterious effects of stress, including a learned aversion to a caretaker^28,35^

A recent work has shown that early-life adversity can alter infant HPA axis reactivity and compromise the mother’s ability to alter pups’ stress hormone response^36^. However, how inherited deficits associated with dysfunctional infant stress systems impact the mother’s ability to regulate her pups stress hormone levels has yet to be explored. Dysfunction of pups’ stress system, especially related to elevated corticosterone levels and its suppression by the mother is of substantial importance because even one day of increased stress hormone levels may have an enduring impact on later life neurobehavioral function^37^.

Here, we examine the ontogeny of threat learning in bLR rats and compare it to the ontogeny of threat learning in outbred SD rats. We also examine whether maternal presence is capable of regulating threat learning and the response to stress in these bLR rats and whether the administration of FGF2 early in life influences the emergence of threat learning in bLR pups.

## Methods

### Animals

Outbred SD (Charles River) breeding pairs and bLR breeding pairs were mated and all pups were born and raised in our colony. The room was set to a 12:12 h light:dark cycle and food and water were freely available. bLR animals were obtained from the Akil laboratory colony. These animals were initially bred from Sprague-Dawley rats obtained from multiple commercial vendors and henceforth selectively bred in the Akil laboratory. bLR pups were offspring of the F43, F48, F50, or F56 generations^9^. All animal handling and behavioral experiments were conducted by a female experimenter. All procedures were approved by the Institutional Animal Care and Use Committee of the University of Michigan.

### Threat Conditioning and Freezing Test

On PND 4, SD and bLR pups were placed in individual plastic beakers fixed with individual Plexon tubing attached to an olfactometer. A heating pad was placed underneath the beakers in order to maintain pups’ body temperature. After 10 minutes of habituation, pups were exposed to 11 30s peppermint odor conditioned stimulus (CS)-1s 0.5 mA shock unconditioned stimulus (US) pairings, 11 unpaired CS and US presentations, or 11 CS presentations. Shocks were manually delivered by the experimenter to the pup’s tail. Each presentation was separated by a 4-minute inter-trial interval^33^. Pups were immediately returned to the home cage at the end of the conditioning session. Beakers were cleaned with water between groups to prevent additional cleaning odors and to prevent the transfer of threat via smell to other experimental groups.

At PND 11, threat memory was assessed by exposing pups to 3 30s CS presentations separated by a 2-minute inter-trial interval. Freezing behavior during the CS presentations was manually scored. Freezing behavior was defined as a clear discontinuation of any movement that is not associated with breathing or other involuntary actions.

### Corticosterone

Pups were threat conditioned or exposed to the odor CS as described above on PND 4 and sacrificed immediately following the end of threat conditioning. A separate group of animals were removed from the home cage and sacrificed immediately in order to establish baseline corticosterone levels. Collection took place between 10am and 2pm. Trunk blood was collected in EDTA tubes which were subsequently centrifuged at 3000 rpm for 10 minutes at 4 C. Following centrifugation, serum was extracted and stored at −80 C until being sent to the UM Core Facility for radioimmunoassay. Radioimmunoassay was performed using the MP Bio Corticosterone Double Antibody RIA Kit (Irvine, CA, USA).

### Maternal Buffering

Dams of the experimental animals or dams of an equivalent postpartum age maintained on an identical diet were deeply anesthetized with sodium pentobarbital prior to maternal buffering experiments. When the toe pinch response was no longer observed, dams were then placed with the experimental animals in an empty cage lined with an absorbent blue pad. Threat conditioning then commenced as described above^33^.

### FGF2

FGF2 (50 ng/g, Sigma) was dissolved in a vehicle solution of 0.1 M PBS with 0.1% BSA. On PND1, bLR pups were injected with FGF2 (20 ng/g, s.c. in 50 ul 0.1 M PBS with 0.1% BSA) or a vehicle solution (s.c. 50 ul 0.1 M PBS with 0.1% BSA) on PND 1^21^. Pups were threat conditioned as described above on PND 4 and either returned to the home cage to undergo a freezing test at PND 11 or were sacrificed immediately after conditioning for corticosterone assay.

### Statistics

Data were analyzed in GraphPad Prism (San Diego, CA, USA) using analyses of variance. Dunnett’s multiple comparisons test and Tukey’s post hoc test were additionally used where appropriate. Differences between groups were considered significant where *p* < 0.05.

## Results

### bLR Pups Show an Early Emergence of Threat Learning and Altered Stress Responsivity During Threat Learning

Outbred SD rat pups that underwent threat conditioning at PND 4 (*n* = 10) did not freeze more to the CS at test than outbred SD rat pups that were exposed to the CS alone (*n* = 9) or the unpaired CS and US (*n* = 8), *F*(2,24) = 0.89, *p* = 0.42 (Figure 1A). However, a 1-way ANOVA comparing mean freezing that bLR pups showed to the CS at test was significant, *F*(2,27) = 5.34, *p* = 0.01 (Figure 1B). Dunnett’s multiple comparisons test revealed that bLR pups that underwent threat conditioning (*n* = 11) froze significantly more to the CS at test than pups that were exposed to the CS alone (*n* = 9; *p* = 0.01) or the unpaired CS and US (*n* = 10; *p* = 0.04).

**Figure 1.**
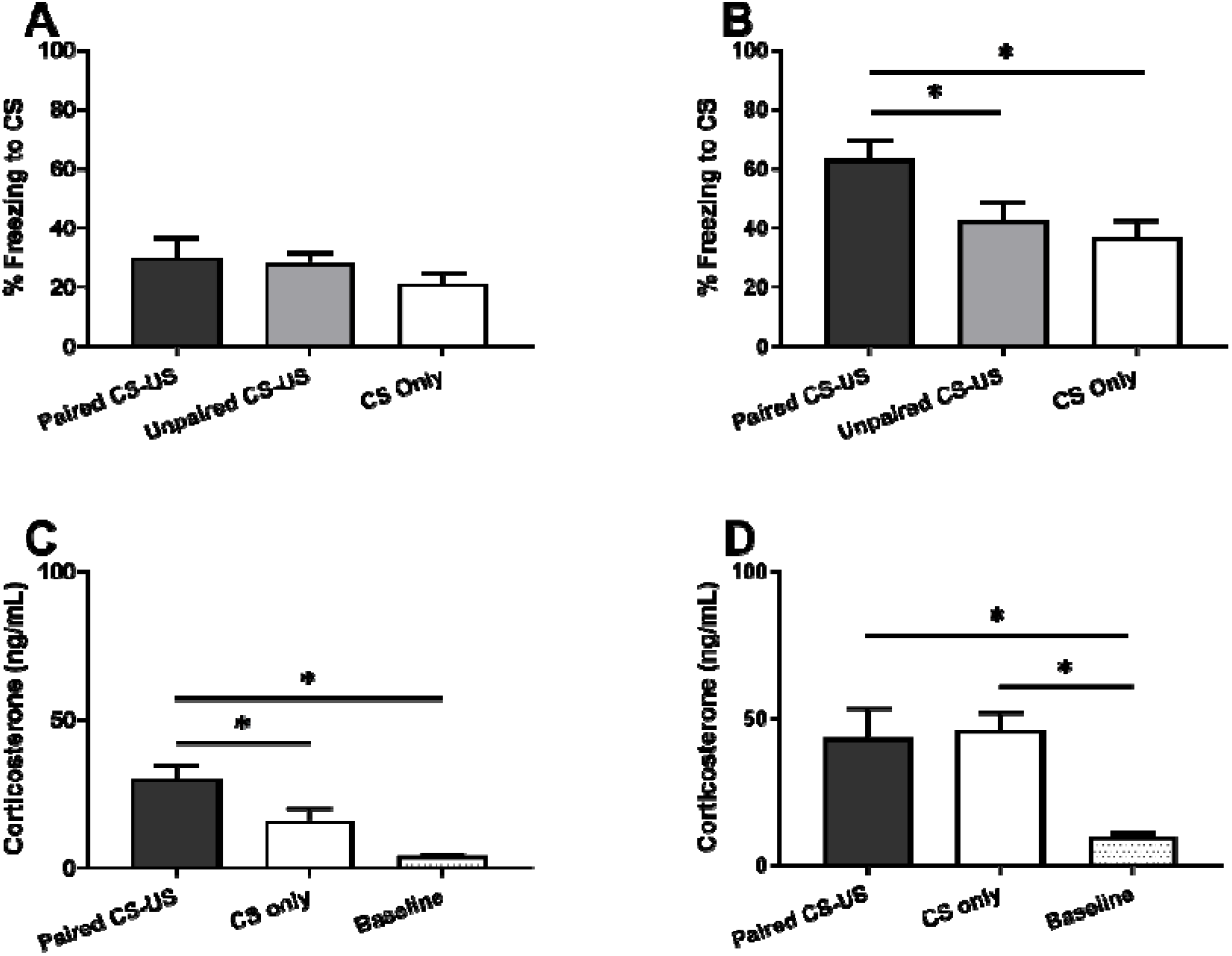
Bred Low-Responder (bLR) Pups Show Early Acquisition of Threat Learning. A. Outbred Sprague-Daw ey pups were exposed to 11 30s peppermint odor conditioned stimulus (CS)-1s 0.5 mA shock unconditioned stimulus (US) pairings. 11 unpaired CS and US presentations, or 11 CS presentations during training a: postnatal day 4, an age at which pups typically do not acquire classical threat learning. Average cue induced freezing was measured during 3 odor CS presentations at postnatal day 11.
B. A: postnatal day 4, bLR pups were exposed to 11 30s peppermint odor conditioned stimulus (CS)-1s 0.5 mA shock unconditioned stimulus (US) pairings, 11 unpaired CS and US presentations, or 11 CS presentations. Average cue-induced freezing was measured during 3 odor CS presentations at postnatal day 11.
C. Average serum corticosterone evets in Outbred Sprague-Daw ey pups exposed to 11 CS-US pairings or 11 CS presentations relative to average serum corticosterone levels from pups taken directly from the home cage.
D. Average serum corticosterone evets In bLR pups exposed to 11 CS-JS pairings. 11 CS presentations.1 CS-US pairings in the presence of an anestheized mother, or taker directly from the home cage.

We next examined the corticosterone response to the experience of threat conditioning or exposure to the CS only at PND 4 in outbred SD rat pups and bLR pups. In outbred SD rat pups, serum corticosterone levels differed across all 3 collection groups, *F*(2,24) = 11.83, *p* = 0.0003 (Figure 1C). Tukey’s post hoc test was used to test for differences in corticosterone levels between each pair of experimental conditions. Pups that underwent threat conditioning showed significantly higher serum corticosterone levels than pups that were exposed to the CS alone (*p* = 0.03) or pups that were taken directly from their cage (*p* = 0.0002). There were no differences in serum corticosterone levels between pups that were exposed to the CS alone or were taken directly from the cage (*p* = 0.10).

In bLR pups, serum corticosterone levels differed across the 3 collection groups, *F*(2,26) = 8.65, *p* = 0.001 (Figure 1D). Tukey’s multiple comparison’s test revealed that relative to corticosterone levels at baseline (*n* = 10), serum corticosterone levels in pups that underwent threat conditioning (*n* = 10; *p* = 0.005) and pups that were exposed to the CS alone (*n* = 9; *p* = 0.003) were significantly higher. There were no significant differences in serum corticosterone levels between pups that underwent threat conditioning and pups that were exposed to the CS alone (*p* = 0.95).

### Maternal Presence During Acquisition Regulates Retention of Learned Threat Responses in bLR Pups But Not Corticosterone Release During Threat Learning

We next examined whether maternal presence was capable of regulating expression of learned threat and corticosterone release during threat learning in bLR pups. These freezing data were compared with the previous freezing data from pups that underwent threat conditioning in the absence of an anesthetized mother. Average freezing to the 3 CS presentations differed across all 4 conditioning groups, *F*(3,36) = 8.74, *p* = 0.0002. Using Dunnett’s multiple comparison’s test, we found that bLR pups that underwent threat conditioning in the presence of an anesthetized mother (*n* = 10) froze significantly less to the CS at test than pups that underwent threat conditioning in the absence of an anesthetized mother (*p* < 0.0001) (Figure 2A).

**Figure 2.**
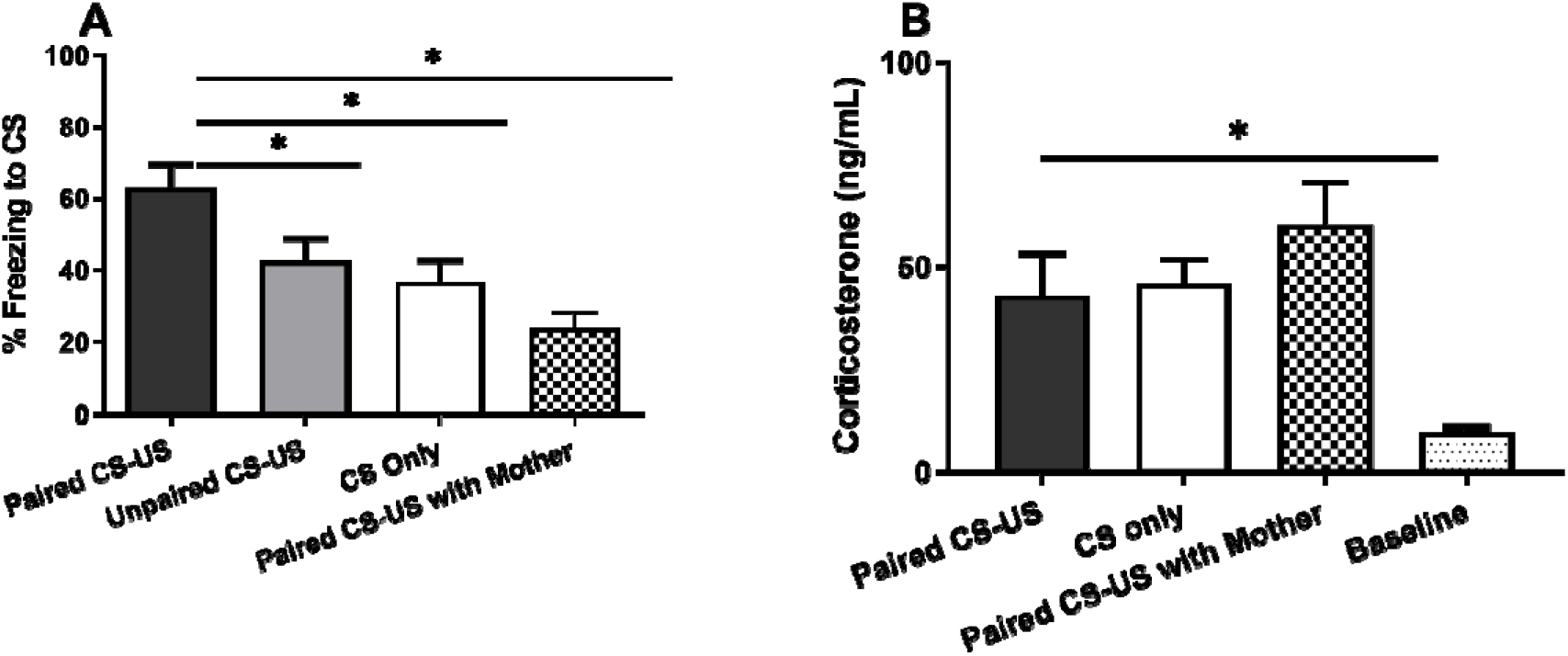
Maternal Presence During Threat Conditioning Can Block Early Expression of Threat Learning In bLR Pups But Not corticosterone Release. A. At Postanatal day 4, bLR pups were exposed to 11 30s peppermint odor conditioned stimulus (CS)-1a 0.5 mA shock unconditioned stimulus (US) pairings, 11 unpaired CS and US presentations, 11 CS presentations, or 11 CS-US pairings in the presence of an anestheized mother. Average cue-induced freezing was measured during 3 odor CS presentations at postnatal day 11.
B. Average Serum corticosterone levels In bLR pups exposed to 1 CS-US pairings, 11 CS presentations, 11 CS-US pairings in the presence of an anestheized mother, or taken directly from the home cage.

To examine whether the mother was capable of regulating corticosterone release during threat conditioning, we collected trunk blood samples from an additional group of bLR pups that were conditioned in the presence of an anesthetized mother. Serum corticosterone levels in these pups were then compared to those from bLR pups in the previous corticosterone experiment. Serum corticosterone levels differed across the 4 collection groups, *F*(3,35) = 7.38, *p* = 0.0006. As described above, serum corticosterone levels in threat conditioned pups and pups that were exposed to the CS alone were significantly higher than baseline levels, *p*’s = 0.02. However, Tukey’s post hoc test found no significant differences in serum corticosterone levels between pups that underwent threat conditioning in isolation relative to pups that underwent threat conditioning in the presence of an anesthetized mother (*n* = 10), *p* = 0.42 (Figure 2B). Serum corticosterone levels were also significantly higher in pups that underwent threat conditioning with maternal presence relative to baseline, *p* = 0.0004.

### FGF2 Administration at PND 1 Blocks the Early Emergence of Threat Learning in bLR Pups but Does Not Modulate Corticosterone Response During Threat Conditioning

Finally, we examined whether early life administration of FGF2 modulated the early emergence of threat learning in bLR pups. At PND 4, threat conditioning or exposure to 11 CS presentations occurred as described above. When exposed to the CS again in a freezing test at PND 11, we observed that FGF2-treated pups (*n* = 16) froze less to the cue than Vehicle-treated pups (*n* = 17), *t*(31) = 2.54, *p* = 0.02 (Figure 3A).

**Figure 3.**
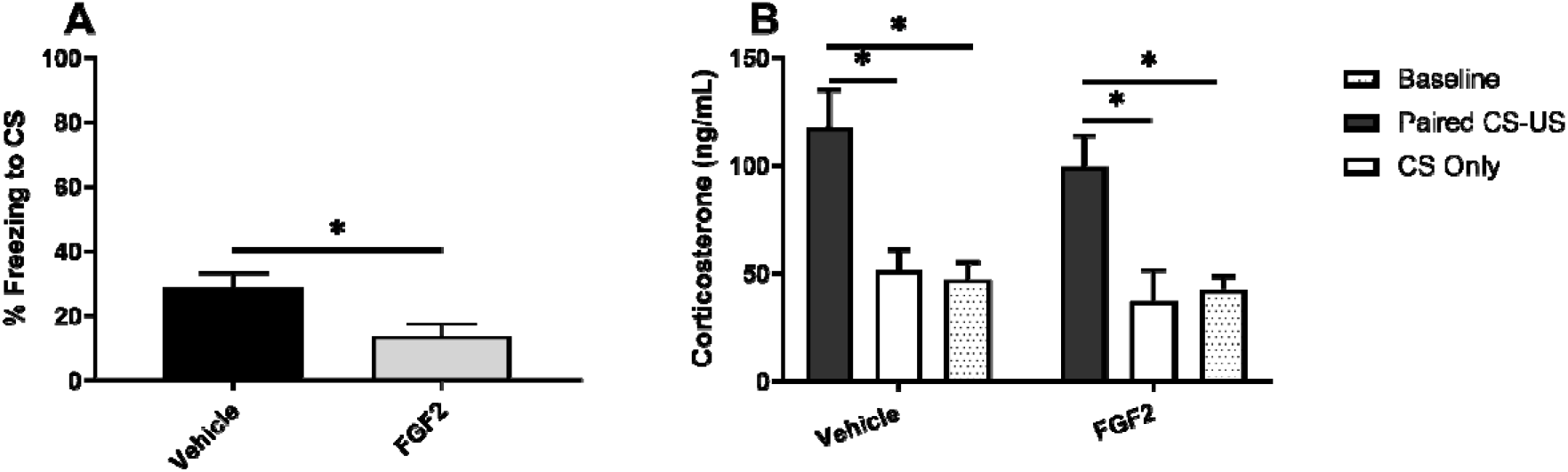
FGF2 Administration on Postnatal Day 1 Can Block Early Expression of Threat Learning in bLR Pups But Not corticosterone Release. A. On postnatal day 4, FGF2 treated or vehicle treated bLR pups were exposed to 11 30s peppermint odor conditioned stimulus (CS) 1s 0.5 tA shock unconditioned stimulus (US) pairings or 11 CS presentations. Average cue-induced freezing was measured during 3 odor CS presentations at postnatal day 1.
B. Average serum corticosterone levels in FGF2-treated or vehicle-treated bLR pups exposed to 11 CS-US parings. 11 CS presentations, or taken directly from the home cage.

In a separate group of animals, we next asked whether early life administration of FGF2 modulated the corticosterone response during threat conditioning. A two-way ANOVA revealed a significant main effect of conditioning group on serum corticosterone levels, *F*(2,33) = 18.14, *p* < 0.0001. However, the effect of FGF2 treatment was not significant, *F*(1,33) = 1.64, *p* = 0.21, nor was there a significant interaction effect, *F*(2,33) = 0.19, *p* = 0.82. Within FGF2-treated pups and vehicle-treated pups, serum corticosterone levels were significantly higher in pups that underwent threat conditioning relative to those that received exposure to the CS alone (*p* = 0.009; *p* = 0.01) and relative to those that were sacrificed immediately after removal from the home cage (*p* = 0.006; *p* = 0.006) (Figure 3B).

## Discussion

Here, we examined the ontogeny of threat learning in a line of rats that show high anxiety-like behavior as a result of selective breeding for inhibited behavior in a novel environment. Previous studies have shown that rat pups younger than PND 10 typically do not show avoidance of odors that have been paired with mild shock^29^ and that freezing behavior in response to noxious stimuli emerges as rat pups approach weaning age^38–43^. Consistent with these findings, we observed that typically-developing outbred SD rats did not show conditioned freezing responses when trained at PND 4 and tested at PND 11. However, we found that bLR rat pups can acquire threat learning and express that learning on a freezing test at PND 11. This early emergence of cued threat-induced freezing could be diminished in bLR pups that underwent threat conditioning in the presence of an anesthetized mother and in bLR pups that received an FGF2 injection on PND 1.

Although there is a significant body of literature supporting the relationship between FGF2 and threat learning, the nature of this relationship differs based on the context. Administration of FGF2 prior to threat conditioning has been shown to enhance contextual threat learning in 19-day old rats^44^. Furthermore, daily administration of FGF2 from PND 1-5 (but not a single administration on PND 1) was associated with a precocious emergence of contextual threat learning on PND 16^45^. Acute FGF2 administration has also been found to facilitate the acquisition of extinction learning and retention of extinction learning^24,25^. Taken together, these results suggest that acute FGF2 may enhance learning processes in general and chronic administration of FGF2 in early life may accelerate the development of contextual threat learning. In bLR animals, a single administration of FGF2 the day after birth suppressed the early emergence of cued threat learning. It is important to note that the above-mentioned studies were performed in wild-type animals, and that here we administered FGF2 to rats of the bLR phenotype. Additionally, the effects of FGF2 on the early emergence of contextual threat learning were observed after multiple FGF2 injections over several days^45^. Other studies have suggested that the effects of early life FGF2 administration are indeed dependent on phenotype^46^.

Although maternal presence during threat conditioning and PND 1 FGF2 treatment were associated with suppressed cue-induced freezing in bLR pups, these manipulations were not sufficient to suppress serum corticosterone levels during threat conditioning. These results are particularly surprising because in typically developing animals, corticosterone appears to regulate the emergence of threat responses. Administration of corticosterone to preweanling pups can drive early expression of a freezing response to predator odor^31^ and the acquisition of threat learning in the presence of the mother^33^. In bLR pups, stress reactivity and threat learning may be operating independently at this age because elevated corticosterone levels did not appear to be sufficient in supporting the acquisition of threat learning in bLR pups. Maternal buffering and FGF2 administration may instead lead to suppressed threat learning by altering neuronal activity within the network of brain areas that underlie infant threat learning or that support the conditioned freezing response^47,48^.

In contrast to what we observed in non-injected bLR pups, we found significantly elevated corticosterone levels in both FGF2 and vehicle-injected bLR pups following threat conditioning relative to following exposure to the CS alone. Baseline levels of corticosterone were also higher in injected bLR pups (regardless of injection type) relative to non-injected bLR pups. The experience of receiving a painful injection in the absence of the dam on PND 1 could have altered the developmental trajectory of bLR pups’ stress response system. Although rat pups are considered to be in a stress hyporesponsive period until about PND 10, studies of typically developing animals suggest that the experience of stress or pain in the absence of the dam can induce acute corticosterone release. Several studies suggest that newborn rats that experience pain in the absence of the dam show elevated corticosterone levels between 24 hours and up to 7 days after the painful experience^49–51^.

The bLR behavioral phenotype appears to be characterized by more robust learned threat responses in infancy in addition to increased spontaneous anxiety-like behaviors observed later in life. Our model of early threat learning in anxiety-like phenotype demonstrates that the relationship between threat conditioning, stress response, and maternal regulation of threat conditioning may be more complex than what studies of typically developing animals show. Our results suggest that although an anxious-like temperament may be associated with early threat learning, environmental factors (such as maternal presence) and pharmacological intervention (such as modulation of the FGF2 system) may be capable of counteracting that early threat learning. Psychosocial and pharmacological interventions in vulnerable infants may therefore increase resilience to adverse events and potentially decrease the risk of developing anxiety disorders later in life.

## Acknowledgements

This research was supported by the K08 MH014743-01A1 and NARSAD Young Investigator Grant from the Brain & Behavior Research Foundation to JD and NIDA U01DA043098; NIH R01MH104261; Office of Naval Research (ONR) N00014-12-1-0366 and 00014-19-1-2149; The Hope for Depression Research Foundation (HDRF): the Pritzker Neuropsychiatric Research Consortium to HA.

